# Maize nested introgression library provides evidence for the involvement of *liguleless1* in resistance to northern leaf blight

**DOI:** 10.1101/818518

**Authors:** Judith M. Kolkman, Josh Strable, Kate Harline, Dallas E. Kroon, Tyr Wiesner-Hanks, Peter J. Bradbury, Rebecca J. Nelson

## Abstract

Plant disease resistance is largely governed by complex genetic architecture. In maize, few disease resistance loci have been characterized. Near-isogenic lines (NILs) are a powerful genetic tool to dissect quantitative trait loci (QTL). We analyzed an introgression library of maize near-isogenic lines (NILs), termed a nested NIL (nNIL) library for resistance to northern leaf blight (NLB) caused by the fungal pathogen *Setosphaeria turcica*. The nNIL library was comprised of 412 BC_5_F_4_ NILs that originated from 18 diverse donor parents and a common recurrent parent, B73. Single nucleotide polymorphisms identified through genotyping by sequencing (GBS) were used to define introgressions and for association analysis. NILs that conferred resistance and susceptibility to NLB were comprised of introgressions that overlapped known NLB QTL. Genome-wide association analysis and stepwise regression further resolved five QTL regions, and implicated several candidate genes, including *Liguleless1* (*Lg1*), a key determinant of leaf architecture in cereals. Two independently-derived mutant alleles of *lg1* inoculated with *S. turcica* showed enhanced susceptibility to NLB. In the maize nested association mapping population, leaf angle was positively correlated with NLB in five recombinant inbred line (RIL) populations, and negatively correlated with NLB in four RIL populations. This study demonstrates the power of a nNIL library combined with high density SNP coverage to resolve QTLs. Furthermore, the role of *lg1* in leaf architecture and in resistance to NLB has important applications in crop improvement.

**Significance Statement (120 words):** Understanding the genetic basis of disease resistance is important for crop improvement. We analyzed response to northern leaf blight (NLB) in a maize population consisting of 412 near-isogenic lines (NILs) derived from 18 diverse donor parents backcrossed to a recurrent parent, B73. NILs were genotyped by sequencing to detect introgressed segments. We identified NILs with greater resistance or susceptibility to NLB than B73. Genome-wide association analysis, coupled with stepwise regression, identified 5 candidate loci for NLB resistance, including the *liguleless1* gene. The LIGULELESS1 transcription factor is critical in development of the leaf ligular region and influences leaf angle. We found that *liguleless1* mutants are significantly more susceptible to NLB, uncovering a pleiotropic role for *liguleless1* in development and disease resistance.

## Introduction

Quantitative trait locus (QTL) mapping has been used to dissect the genetic architecture of many important agronomic traits in crop plants, including disease resistance. Most disease resistance QTLs have small to moderate effects on disease phenotypes, and the underlying polymoprhisms are not known. Quantitative disease resistance loci have typically been identified through associations between DNA markers and disease phenotypes through linkage mapping or genome wide association studies (GWAS). The maize nested mapping association mapping (NAM) population (Yu et al., 2008), a multi-parent recombinant inbred line (RIL) population, has increased the resolution of QTLs for many traits, including several disease traits (Kump et al., 2011; Poland et al., 2011, Benson et al., 2015, Li et al., 2018). A population of near-isogenic lines (NILs) was developed in parallel with the NAM (Gandhi et al., 2008), and has been used for detailed analysis of selected QTL (Ding et al., 2017).

Comparing a NIL to the corresponding recurrent parent allows the effects of specific chromosomal segment(s) to be assessed. The uniform genetic background allows detection of smaller allelic effects in NILs relative to RILs (Keurentjes et al., 2007; Marchadier et al., 2019). The use of NILs can permit diverse alleles at a locus of interest to be compared in a common and adapted genetic background, potentially allowing the identification of novel sources of variation for breeding programs (Young et al., 1988; Bernacchi et al., 1998). QTL can be resolved to detect candidate or causal genes through breakpoint analysis by further backcrossing of selected NILs for fine mapping (Eshed and Zamir, 1995; Monteforte and Tanksley, 2000; Jeuken and Lindhout, 2004). Utilizing RILs typically allows QTL mapping at low resolution, and fine mapping with NILs can also be limited in regions of low recombination (e.g., Jamann et al., 2014). NILs have been used in genetic studies of quantitative disease resistance in maize for QTL discovery (Chung et al., 2010; Liu et al., 2015; Lopez-Zuniga et al., 2019), for phenotypic analysis (Chung et al., 2011; Jamann et al., 2014; Zhang et al., 2017) and for identifying the genes underlying QTL (Yang et al., 2017).

Resistance to northern leaf blight (NLB), caused by the fungal pathogen *Setosphaeria turcica*, is inherited both quantitatively and qualitatively. Many QTL and several race specific genes for NLB have been identified in biparental mapping populations (Welz and Geiger, 2000; Wisser et al. 2006; Galiano-Carneiro and Miedaner, 2017). The genetic architecture of NLB resistance was further analyzed in the NAM population (Poland et al., 2011), which consists of 5,000 RILs derived from crosses between B73 and 25 diverse founder lines (Buckler et al., 2009, McMullen et al., 2009, Yu et al., 2008). Using the NAM population, 23 and 49 NLB QTL were identified using 1,106 SNPs (Poland et al., 2011) and 7,386 SNPs (Li et al, 2018), respectively. Several NILs carrying major and minor NLB loci have been developed and characterized for resistance to diseases (Chung et al., 2010; Chung et al., 2011; Balint-Kurti et al., 2010; Belcher et al., 2012; Lopez-Zuniga et al., 2019). Many NLB QTL, however, remain to be characterized in detail.

This paper presents the characterization of a set of ∼400 maize NILs derived from a subset of the maize NAM founder (donor) lines backcrossed to the recurrent B73 inbred line. We used genotyping by sequencing (GBS; Elshire et al., 2011) to provide high density SNP coverage across the collection of NILs, termed here as a ‘nested NIL’ (nNIL) library. The nNIL library was analyzed for resistance to NLB through three years of inoculated field trials. Because the donor lines are genetically diverse, both the ancient and recent recombination events within the uniform genetic background were harnessed using genome wide association studies (GWAS) to aid in resolving QTL for resistance to NLB. While not all QTL were represented by a sufficient diversity of donor lines or NIL coverage to allow extensive QTL dissection through association analysis, five regions of the genome harboring NLB QTL were targeted with significant resolution. In one such region, the *liguleless1* (*lg1*) gene was implicated through GWAS. Inoculation of lines carrying mutant alleles of *lg1* showed that *lg1* importantly plays a previously undescribed role in resistance to NLB, suggesting that *lg1*, in addition to regulating leaf architecture traits, may control factors that influence disease resistance.

## Results

### Characterization of the NILs

We analyzed a set of NILs that were derived from crosses between 18 diverse maize inbred lines and the recurrent parent B73 (termed nNILs for the nesting of diverse haplotypes within introgressed intervals). The donor lines were founders of the widely-utilized NAM population (Fig. S1; Buckler et al., 2009, McMullen et al., 2009, Yu et al., 2008). In total, we identified 992 introgressions across 412 NILs, covering the entire genome (Fig. 1; Table S1). There were on average 2.4 introgressions per NIL, with 29% (n=120) of the NILs having only one introgression. The total set of introgressions summed to 24.3 Gb, with an average of 59.0 Mb per NIL, or 2.9% of the genome per NIL. The average introgression size across the genome was 24.3 Mb with a median introgression length of 9.3 Mb, and a range from 59 to 188,059 Kb. Over 52% of the introgressions were under 10 Mb. In general, longer introgressions traversed the centromeres, and shorter introgressions were identified near the ends of the chromosomes (Fig. 1). The size of the introgressions sharply increased with the distance from the chromosome ends. Introgressed regions spanned the genome and did not appear to be biased towards a given genomic region or introgression donor (Fig. S2). The 992 introgressions were comprised of 621 homozygous introgressions, 212 heterozygous introgressions, and 159 introgressions that included both homozygous and heterozygous segments (Fig. S3). The 992 introgressions included 840 segments that were homozygous and 417 segments that were heterozygous. The introgressed segments were 81% homozygous for a total of 19,703 Mb (average = 23.5 Mb), and 19% heterozygous, totaling 4,608 Mb (average = 11.0 Mb).

**Figure 1.**
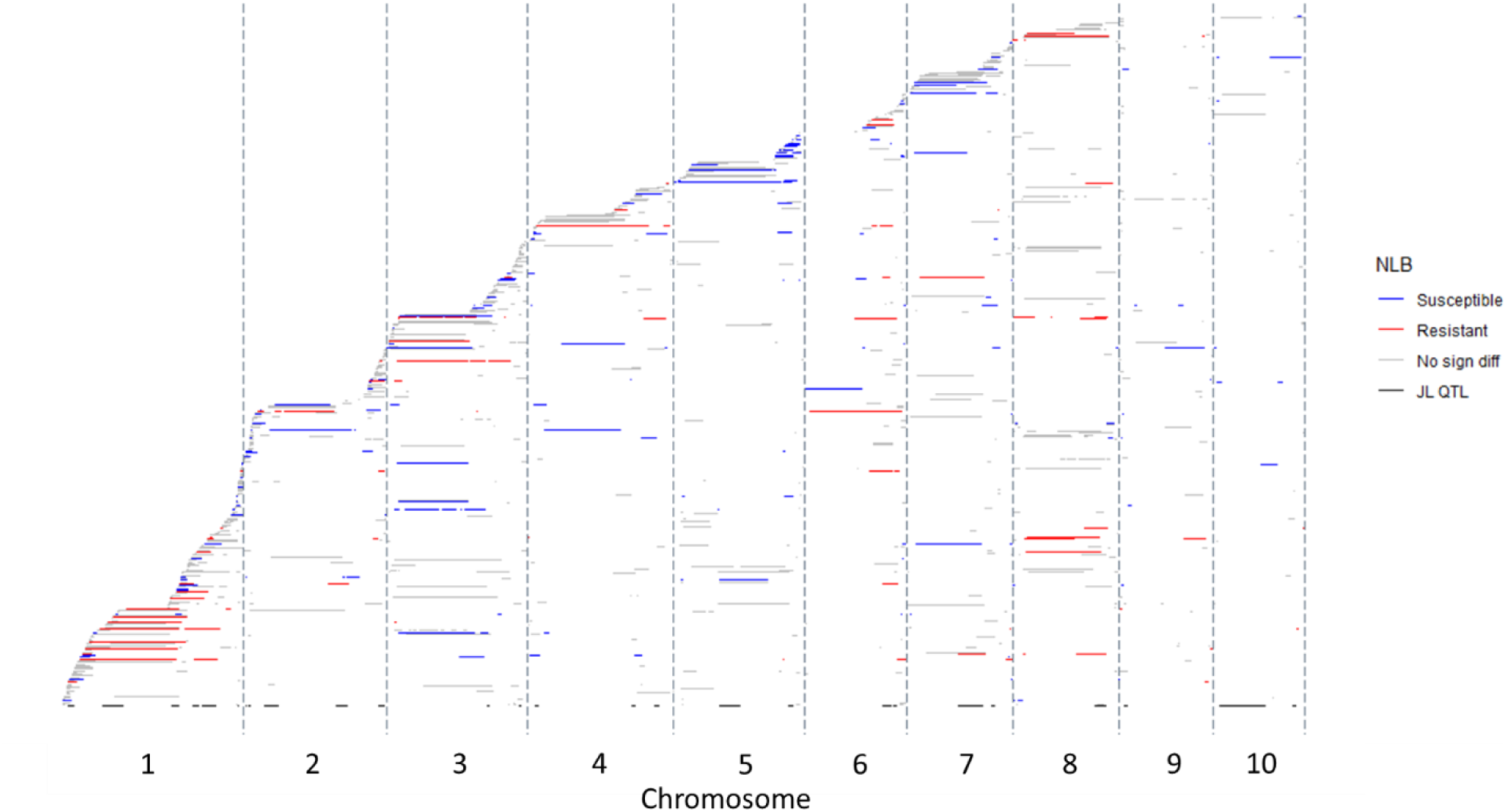
Introgressions in near-isogenic lines (NILs) across the maize genome, ordered along the y-axis based on the first introgression start site. Introgressions marked in red, blue and gray represent NILs that were more resistant, susceptible or not significantly different to northern leaf blight (NLB) in comparison to B73, respectively. Black bars along the X-axis represent the NLB quantitative trait loci mapped in the nested association mapping population (Poland et al., 2011). Grey vertical bars delineate the chromosomes.

Across the majority of NILs, two regions of the genome harbored what appeared to be introgressions, but were more likely sequencing artifacts or polymorphisms associated with different B73 seed sources used in different research groups (Liang and Schnable, 2016). Between the regions of approximately 289.85 Mb and 293.15 Mb of chromosome 1, SNP variants appeared to produce two main haplotypes across the set of NILs. A second region with SNP variants across the entire NIL set was identified on chromosome 5, between approximately 208.16 and 211.54 Mb; this region also showed two main haplotypes. These sequence artifacts were not included in the defined introgressions.

### Phenotypic traits in the nNIL library

Phenotypic analysis of the nested NIL library, consisting of 453 NILs, was conducted in 2011-13 for days to anthesis (DTA) and three times post-flowering for diseased leaf area (DLA). The average (and range) of DLA for 2011, 2012 and 2013 was 27% (6.3-50%), 26% (5.3-52%) and 26% (6.0-54%), respectively, with B73 averaging 30% (2011), 25% (2012), and 21% (2013). Area under the disease progress curve (AUDPC) and DTA across years appeared to be normally distributed Fig. S1). Genotype and genotype-by-environment effects were significant for AUPDC, while replication and environment effects were not. The estimated heritability on an entry means basis was 0.74 for AUDPC and 0.36 for DTA. The average (and range) of DTA across the three years was 68.4 days (61.0-80.0 days), 76.8 days (68.5-84.5 days) and 88.0 days (81.0-103.0 days) respectively, with the average DTA of B73 being 68.4 days (2011), 76.8 days (2012) and 86.3 days (2013). The genotype effect was significant for DTA while the environment, replication and the genotype by environment interaction were not. In the nNIL library, there was a low but significant negative correlation between AUDPC and DTA (*R*^*2*^=- 0.025; *p*=0.0009; Fig. S1). In contrast, there was a highly significant negative correlation between AUDPC and DTA in the 282 maize diversity panel (*R*^*2*^ = −0.39; *p*<0.0001; Fig. S4), which may largely be due to population structure. All but the tropical subpopulations showed a significant negative correlation between DTA and AUDPC. In general, there was less variability in NLB and DTA in the tropical subpopulation, which as a group was more resistant to NLB, but also had greater DTA in comparison to other subpopulations.

Estimated allelic effects for resistance to NLB (calculated as AUDPC, based on % DLA) in the NILs ranged from −13.7 to 16.0 (mean = 0.00). In contrast, the range of allelic effects for resistance to NLB (based on % DLA) in the NAM Joint Linkage QTL (Poland et al., 2011) ranged from −5.6 to 5.3 (mean = 0.07). Four NILs showed significantly higher levels of NLB resistance relative to B73 based on the Dunnett’s multiple test comparison. An additional 33 NILs were more resistant to NLB than B73 based on a 95% confidence interval (Table S2). The most resistant NIL had a large introgression on chromosome 6, covering a known QTL at bin 6.05 (Chung et al., 2011; Poland et al., 2011). Averaged across 3 years, this line showed a 17.5% decrease in NLB. The other three most resistant NILs based on Dunnett’s test for multiple comparison all harbored introgressions covering a large section of chromosome 1 and showed 15.6, 14.7 and 13.9% lower NLB than B73. Twenty NILs were significantly more susceptible to NLB than B73 based on the Dunnett’s multiple test comparison, with 56 additional NILs more susceptible to NLB based on a 95% confidence interval.

Thirty-four NILs had significantly higher DTA relative to B73, and three NILs had significantly lower DTA relative to B73 based on a Dunnett’s test. Five of the 34 NILs also had significantly different to DTA relative to B73 (Table S2). Three of the NILs with an effect on DTA had been identified as more susceptible to NLB (compared to B73). These included a NIL with an introgression on the end of chromosome 1; a NIL with a large introgression on chromosome 8 that overlaps with flowering time locus *Vegetative to generative transition1* (*Vgt1;* (Salvi et al., 2007; Ducroq et al., 2008) and two other introgressions; and an NLB-susceptible NIL with an introgression at 12-36 Mb on chromosome 1, and second small introgression (<2 Mb) on chromosome 7. Two NILs with significantly different DTA were more resistant to NLB. The NIL with the largest days to anthesis harbored an introgression that spans a large area on chromosome 8 where both major flowering time (*Vgt1*) and NLB resistance *genes Helminthosporium turcicum resistance2* (*Ht2)* and *Helminthosporium turcicum resistanceN1* (*HtN)* reside (Chung et al., 2010; Hurni et al., 2015). The other NLB-resistant, high-DTA NIL had a small introgression on chromosome 1, and large introgressions on chromosomes 7 and 8 (Tables S1 and S2).

The introgression tiling path revealed several trends (Fig. 1). Introgressions from any single parent were distributed across the genome (Suppl Fig. 3), making breakpoint analysis from individual donors an unsuitable means to resolve QTLs. As expected based on patterns of recombination (Rogers-Melnick et al., 2015), smaller introgressions were identified on the ends of the chromosomes (Fig. 1). Introgressions spanning centromeres tended to be large, and two regions of the genome were associated with particularly large, resistance-associated introgressions. Introgressions spanning the centromeric region of chromosome 1 (∼80 to 200 Mb) were frequently associated with resistance to NLB, perhaps reflecting the known NLB QTL in bin 1.06 (Wisser et al., 2006; Chung et al., 2011; Jamann et al., 2014). Introgressions on chromosome 8 also tended to be large and associated with resistance, likely reflecting the major genes and QTLs that have been identified in that region (Chung et al., 2010; Wisser et al., 2006; Hurni et al., 2015). In contrast, introgressions on chromosome 5 were often associated with increased susceptibility to NLB compared with B73, which has a known NLB resistance QTL in bin 5.05 (Poland et al., 2011; Li et al., 2018, Wisser et al., 2006). Although the B73-derived resistance in bin 5.05 was displaced by introgressions that increased susceptibility in seven NILs, breakpoint analysis was not successful in resolving the QTL, which spanned the 9.5 Mb region (Fig. 1).

**Figure 2.**
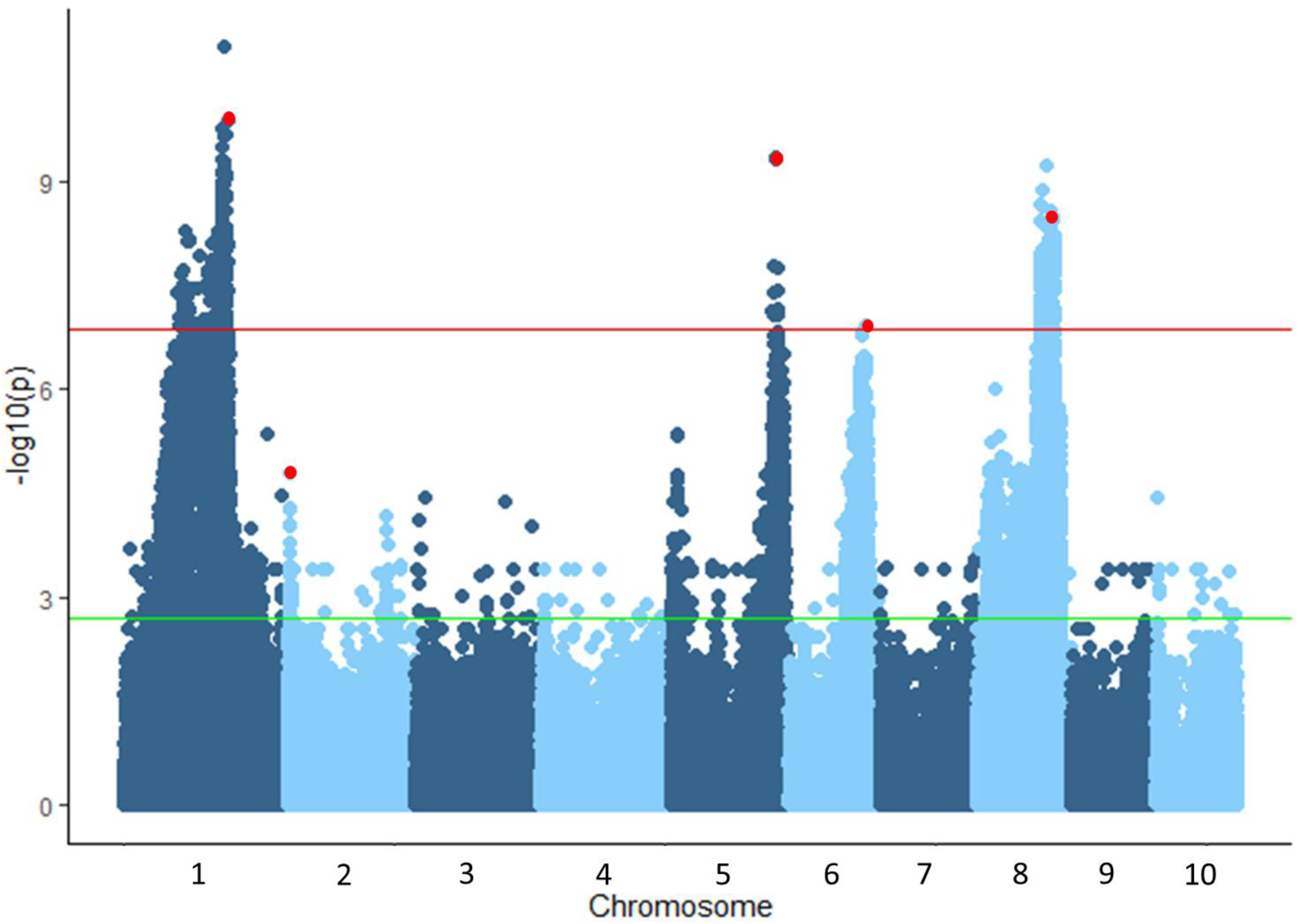
Association analysis of resistance to northern leaf blight in the nested near-isogenic line library using the general linear model in Tassel. Significance thresholds were determined using the Bonferonni correction factor (red line) and the false discovery rate (green line). The SNPs most significantly associated with resistance by stepwise regression are highlighted in red.

**Figure 3.**
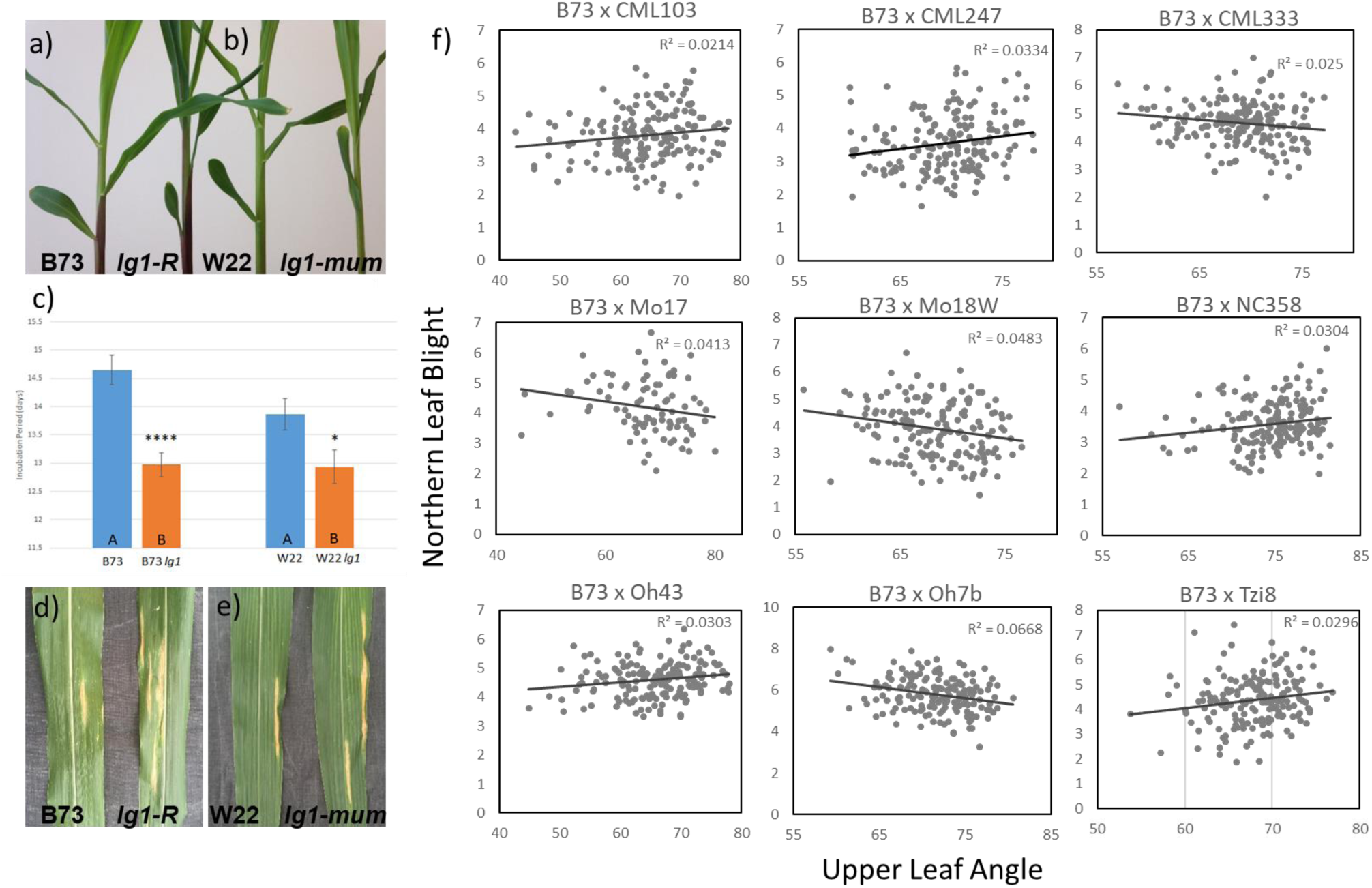
The effect of *liguleless1* (*lg1*) on resistance to northern leaf blight. Leaf architecture is impacted by mutations in *lg1* (a) B73 (left) and *lg1-R* (right), and (b) W22 (left) and *lg1-mum* (right). (c) The *lg1-R* and *lg1-mum* alleles were both significantly more susceptible to northern leaf blight (NLB), measured as incubation period in the greenhouse (the number of days after inoculation that the NLB lesion appeared). NLB lesions on the leaves of (d) B73 and *lg1-R*, and (e) W22 and *lg1-mum*. Significant positive and negative correlations (f) were identified between the upper leaf angle (Tian et al., 2011) and resistance to NLB (NLB index; Poland et al., 2011) in 9 RIL populations of the nested association mapping population.

### Association analysis for AUDPC in the NIL library

GWAS was employed in the nNIL library with 374,145 genome-wide SNPS to take advantage of the polymorphisms among the nNIL parental lines. After Bonferroni correction, four regions of the genome were significantly associated with resistance to NLB (Fig. 2). The four implicated regions overlapped with known NLB QTL on chromosomes 1, 5, 6 and 8 (Poland et al., 2011; Wisser et al., 2006). The top three SNPs in each of these significant peaks were inferred to implicate nearby candidate genes (Table 1).

**Table 1.**
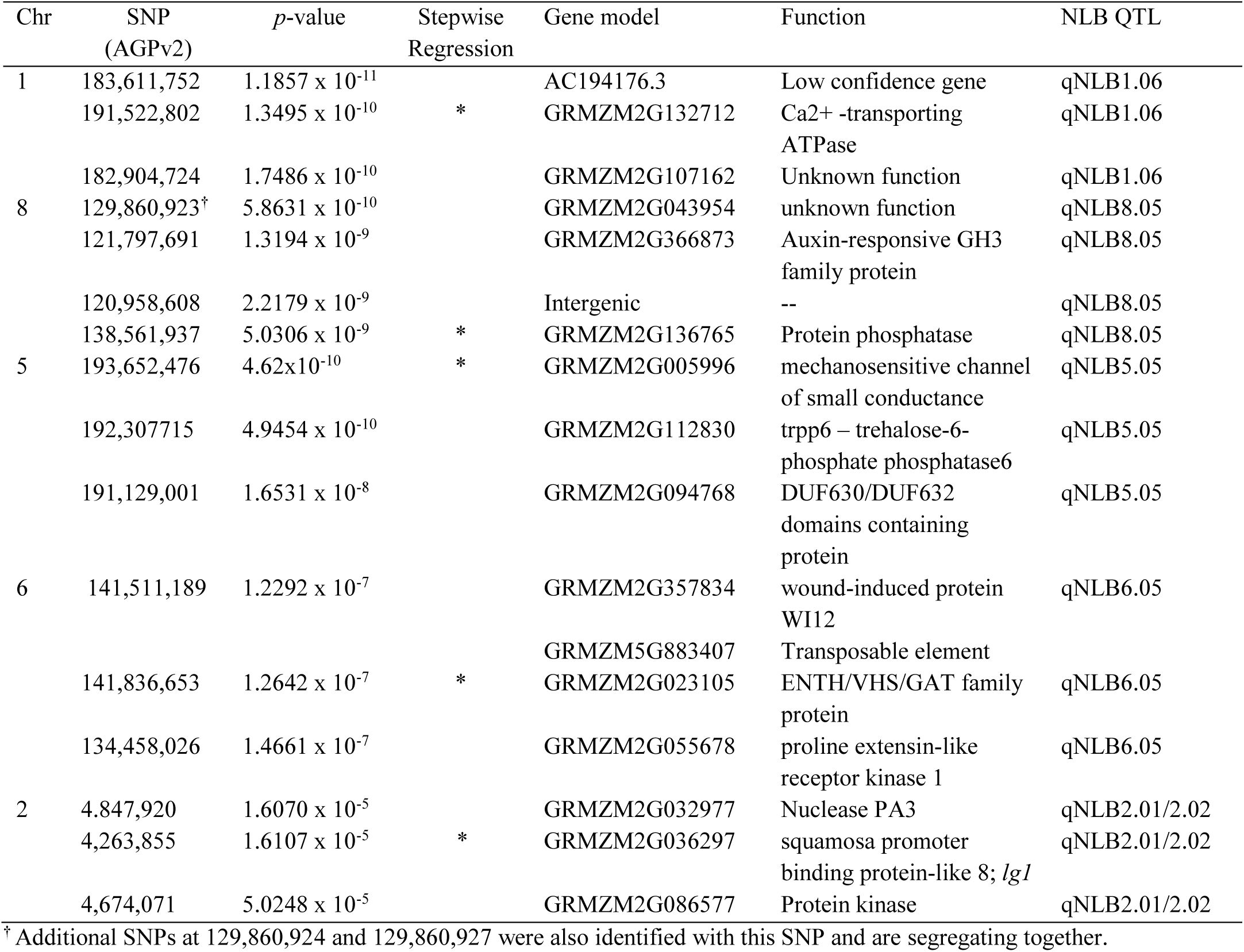
SNPs associated with resistance to northern leaf blight (NLB) in the nested near-isogenic line library were identified through genome-wide association. The top three SNPs are listed along with corresponding attributes, across the 4 regions that were identified based on a Bonferonni threshold. An additional SNP/region was identified via stepwise regression. The 5 SNPs that were identified as contributing to resistance to NLB based on stepwise regression are indicated below with an *.

Stepwise regression further identified regions associated with disease phenotypes. Five significant SNPs were detected, accounting for variation in each peak that was identified based on GWAS as well as an additional SNP on chromosome 2 (Table 1). The large region spanning the centromere of chromosome 1 (80-200 Mb) had a large region with significant association, reflecting the lack of recombination and the consequently large introgressions found in NILs covering this region. The most significant SNP identified by GWAS was in a low confidence gene at 1:183,611,752, while the second most significant GWAS association and the most significant SNP identified through stepwise regression at 1:191,522,802, in a gene annotated as a Ca+ transporter. Using GWAS and stepwise regression, the most significant SNP at the QTL on Chromosome 8 was in a gene identified as a protein phosphatase, located at 8:138,561,937. The significant SNP identified through GWAS and stepwise regression on chromosome 6 was located in a gene annotated as encoding an ENTH/VHS/GAT family protein (Table 1).

B73 was the source of the more resistant allele for two of the NLB QTL regions. Two significant SNPs around the qNLB5.05^NAM^ region were identified via GWAS, one of which was located in a gene (GRMZM2G005996) annotated as a mechanosensitive channel of small conductase. Additionally, this SNP (5:193,652,476) was identified at the same locus via stepwise regression (Table 1). The second SNP (5:191,129,001) identified via GWAS was located in GRMZM2G112830, a gene annotated as trehalose-6-phosphate phosphatase6. The two genes are separated by approximately 1.3 Mb, substantially reducing the candidate interval for qNLB5.05_NAM_ from a previously reported 9.5 Mb (Poland et al., 2009).

The second B73-derived NLB QTL region identified via stepwise regression rested on the short arm of chromosome 2, overlapping with qNLB2.01/2.02_NAM_ (Poland et al., 2011). The identified SNP (2:4,263,855) was one of two peaks identified via GWAS in the QTL, but the only one identified as significantly contributing towards resistance. The SNP (2:4,263,855) was located immediately downstream of the *liguleless1* gene (*lg1,* GRMZM2G036297). The second SNP (2:4.847,920) identified via GWAS was located in GRMZM2G032977, annotated as a nuclease PA3. *Liguleless1* was previously identified as a candidate gene via GWAS in the NAM population (Poland et al., 2011).

### liguleless1 is a candidate gene for resistance to NLB

Mutant analysis was conducted to probe the role of the *lg1* gene in the NLB phenotype. Lines carrying two mutant alleles were examined for NLB response in greenhouse trials: the *lg1-R* allele in the B73 background and the *lg1-mum* allele in the W22 background. Incubation period was measured after whorl inoculation with *S. turcica*, which reduced the influence of leaf angle on disease development. Both mutant alleles were significantly more susceptible to NLB compared to respective inbred lines (Fig. 3). Incubation period was approximately 1.5 days and 1 day shorter in the *lg1* mutant relative to B73 and W22 inbred lines respectively, indicating increased susceptibility through a more rapid infection cycle in the *lg1* mutants.

In maize, mutations in *lg1* result in more upright leaves (Moreno et al., 1997) and natural variation in *lg1* associates with more upright leaves in maize breeding lines (Dzievit et al., 2019; Tian et al., 2011). We tested the relationship between leaf angle and resistance to NLB. A significant negative correlation was identified between middle leaf angle and resistance to NLB (AUDPC) in the 282-line maize diversity panel (*p* < 0.001; *R*^*2*^ = 0.175; Flint-Garcia et al., 2005; Fig. S5)). This association did not account for population structure or kinship. At the sub-population level, significant negative correlations between leaf angle and AUDPC were observed in the non-stiff stalk (NSS) and mixed subpopulations (*p* < 0.0001; *R*^*2*^ = 0.285 and *p* = 0.0006; *R*^*2*^ = 0.106 respectively; Fig. S6). In these sub-populations, the higher leaf angle (more erect leaves) was correlated with lower levels of disease. Leaf angle and resistance to NLB were not significantly correlated within the stiff stalk (SS), tropical, popcorn or sweet corn sub-populations. GWAS utilizing the 282 maize diversity panel, using the mixed linear model (MLM) to account for population structure and kindship, did not identify a significant association with NLB across *lg1* (Fig. S6). To further assess the effect of leaf angle on resistance to NLB, we examined the correlations of upper leaf angle and resistance to NLB in the individual RIL populations of the NAM populations (Tian. et al., 2011; Poland et al., 2011). Of the 26 RIL populations, five exhibited significant positive correlations between leaf angle and NLB, while four exhibited a significant negative correlations between leaf angle and NLB (Fig. 3). The roles of *lg1* in influencing leaf angle and NLB pathogenesis are distinct and putatively multi-allelic.

## Discussion

The genetic architecture of many maize traits have been studied in the NAM population, including flowering time (Buckler et al., 2009); leaf architecture (Tian et al., 2011); plant height (Peiffer et al., 2014); stalk strength (Peiffer et al., 2013); resistance to NLB (Poland et al., 2011), southern leaf blight (Kump et al., 2011), and grey leaf spot (Benson et al., 2015); hypersenstivie defense response (Olukolu et al., 2014); kernel composition (Cook et al., 2012); photosynthesis (Wallace et al., 2014); inflorescence architecture (Brown et al., 2011); and photoperiod sensitivity (Hung et al., 2012). We show the power of using a nested NIL library derived from the parents of the NAM population in the validation, phenotypic characterization, fine-mapping, and gene discovery of the QTL for NLB.

### Description and utility of the nNIL library for dissection of QTL

The nNIL library described in this paper is part of a community resource developed by Syngenta as a tool for dissection of important traits (Gandhi et al., 2008) and available upon request from the International Maize and Wheat Improvement Center (CIMMYT). We describe a subset of a larger population of NILs that was derived from 18 donor parents (consisting of the founder lines in the NAM population) and backcrossed to B73. The potential for association mapping in a nNIL population is based on the diverse parentage of the nNILs and the high-density SNP coverage, as obtained in this study. The library can be used for QTL mapping, and individual NILs can be selected for use in fine-mapping of specific loci of interest, to allow testing of candidate genes. The heterozygous introgressions identified in these lines can be exploited for heterozygous inbred family analysis (Tuinstra et al., 1997).

Recombination rates vary across the genome, with increased recombination near the telomeric regions and decreased recombination near the centromeres (Rodgers-Melnick et al., 2015). Consequently, the introgression sizes identified in this study varied widely and as generally predicted. QTL resolution was poor in regions around centromeres, as expected. Keurentjes et al. (2007) compared QTL resolution in Arabidopsis, and found that, given similar population resources, RILs provided greater QTL resolution than individual NILs at defining QTL intervals. In this study, we utilized approximately 9% of the number of NILs compared to RILs utilized in the NAM population. The average size of NLB QTL in the NAM RILs was 14.6 Mb (Poland et al., 2011), which was smaller than the average introgression size in our NIL library (24.3 Mb), although the range of the introgression sizes was much larger in the nNIL library than compared to that identified through the joint linkage mapping of NLB QTL in the NAM population. The median size of a NAM NLB QTL (9.5 Mb) was, however, much more similar to that of the median NIL introgression size (9.3 Mb).

Across the NILs, the range in introgression size varied, with the smallest introgression in the NIL library (59.2 Kb) being smaller than the smallest NAM NLB QTL (855 Kb), and the largest introgression size (188 Mb) being considerably larger than the largest NLB QTL (77.1 Mb). Thus, while the NIL introgressions are generally larger than the average NAM NLB QTL, the range of introgression size is wider. Because of the SNP saturation across the genome, we were able to detect introgressions smaller than 5 Mb, which might not have been detectable with a sparser marker dataset. These micro-introgressions do not have the characteristics of sequence artifacts, so are provisionally presumed to be valid.

A NIL library in Arabidopsis detected QTL with smaller allelic effects than those in the corresponding RIL population (Keurentjes et al., 2007). In the current study, the mean allelic effect size of QTL detected in the nNIL library was similar to that identified in the NAM RIL population (Poland et al., 2011). However, the range in allelic effects of the nNIL library was approximately three times the size of the allelic effect size of the joint linkage NAM QTL. The homogeneity of the genetic background should make it able to identify smaller effect QTL. We did not identify additional QTL of smaller effect, apart from those with flowering time effect, in comparison to the set identified in the full NAM population, likely due to the smaller sample size used in this study. The nNIL population size tested was approximately 9% that of the NAM RIL population.

We identified NILs that were either more resistant or more susceptible to NLB than B73, implicating loci for resistance to NLB across the genome. The QTL identified were consistent with those identified in the NAM RIL population. QTL with large effect sizes, associated with donor-derived introgressions, were identified across the centromeric regions of chromosomes 1, 6 and 8. These regions overlapped with the previously described QTL designated as qNLB1.06 and qNLB6.05 (Jamann, et al. 2014; Poland et al, 2011). NILs with introgressions across the centromeric region of chromosome 8 overlapped two QTL for resistance to NLB, including qNLB8.06, which overlaps the *Ht2* and *Htn* resistance loci (Chung et al., 2010; Hurni et al., 2015). Further backcrossing, together with inoculation of different *S. turcica* races, will allow development of a “differential series” for major and minor resistance loci for NLB in the B73 background. Inoculation of the lines carrying the 8.06 QTL may allow the specific loci to be identified. Inoculation of other lines will allow the possible race-specificity of QTL to be established. The introgressions in NILs that span qNLB1.06 and/or qNLB8.06 are large, however, suggesting that lack of recombination will inhibit fine-mapping in those regions, as has been previously observed in at qNILB1.06 (Jamann et al., 2014).

B73 is considered moderately susceptible to NLB, and is known to carry alleles for NLB resistance QTL (Poland et al., 2011). In this study, we identified twice as many NILs that were more susceptible to NLB than B73 (lines in which B73-derived resistance was lost with an introgression), than NILs that were more resistant than B73. Additionally, resistance in two of the five QTL regions identified in this study via GWAS and stepwise regression was derived from B73, the recurrent parent of the nNILs: qNLB 2.01 and qNLB5.05. The design of the nNIL library, like the NAM population, was favorable for identifying resistance derived from B73; the nNIL library allowed systematic analysis of the recurrent parent genome for effects on disease, while the donor genomes were only partially represented in the set of nNILs tested here.

A previous study in the 282 maize inbred diversity panel showed a negative genetic correlation between DTA and AUDPC (Wisser et al., 2011) that observed in the sub-populations of the 282 maize inbred diversity panel. In comparison to the other subpopulations, the tropical subpopulation, generally had limited variation for both DTA (showing longer days to flowering) and AUDPC (showing higher levels of resistance). A low negative correlation was detected between DTA and NLB in the nNIL library. This may be due to a small number of NILs with introgressions that were more resistant to NLB and had longer flowering time, either due to linkage (different loci influencing the two traits) or pleiotropy (individual gene[s] influencing both traits). While a significant negative correlation between DTA and resistance to NLB was found in the NAM founder lines, significant negative correlations were found in only 8 of the 26 NAM RIL populations (Poland, 2010).

The lower correlations between maturity and disease resistance in the RIL and NIL populations indicates that some of the correlations seen across germplasm collections may be due to population structure rather than to linkage and/or pleiotropy (Poland et al., 2010). Despite the overall trend, an opposite effect of days to anthesis on NLB was identified. A NIL harboring an introgression that spanned the qNLB1.02 region had increased NLB resistance but reduced days to anthesis relative to B73, confirming a previous report (Jamann et al., 2016). Several NILs with longer maturity harbored introgressions at the telomeric end of chromosome 1 that had not been identified previously in NAM QTL mapping. These introgressions either were identified in our study due to an increase in allele effect sensitivity in our population, or because these introgressions co-localized with flowering time effect, which may be been a reduced factor in the NAM QTL mapping where DTA was used as a covariate.

### Association analysis in the nNIL library utilizes ancient recombination for greater QTL resolution

To identify candidate loci or regions for resistance to NLB, we utilized the polymorphisms among the introgression donors at a given locus using association analysis across the nNILs. The uniform B73 genetic background diminished the confounding effect of population structure and kinship. Only a small number of NILs had introgressions in any given genomic region, yet significant effects were identified at 5 regions in the genome with known NLB QTL. Utilizing high density SNP coverage in the ∼400 NILs, we were able to narrow the interval of interest in a region of low recombination (qNLB1.06), identified putative candidate alleles in two other QTL regions of large effect (qNLB8.05, and qNLB6.05), and provided a narrower window of significance for two QTLs with B73-derived resistance. One of the B73 QTL, qNLB5.05, was narrowed to two significant peaks about 1 Mb apart, which might indicate a multi-gene QTL. The other B73-derived QTL, qNLB2.01/2.02, harbored two SNPs, approximately 0.6 Mb apart, with the most significant SNP being 600bp from the *Liguleless*1 gene

### Candidate gene analysis of qNLB2.01 implicates *liguleless1*

A significant SNP in qNLB2.01/2.02 was identified within ∼600 bp of the *liguleless*1 gene. The resistant allele was derived from B73. NAM GWAS analysis had previously identified a SNP in the *lg*1 gene and within 100 kb of *lg1*, using HapMapv1 (Poland et al., 2011), and HapMapv2 (Chia et al., 2012) respectively. We used controlled inoculation of *lg1* mutant lines to further probe the role of *lg1* in disease resistance. Because the *lg1* gene influences leaf angle, we chose a disease assay that was intended to minimize the influence of microclimate due to leaf angle or canopy architecture: we applied fungal spores in the whorl and scored the time to primary lesion formation. The two mutants tested were significantly more susceptible to NLB than their wild type counterparts, B73 and W22, with a decrease in incubation period of 1.5 and 1 days, respectively.

The natural variant and mutant alleles of *liguleless1* impact leaf angle and subsequent canopy structure, that ultimately affects plant density (Becraft et al., 1990; Sylvester et al., 1990; Tian et al., 2011; Tian et al., 2019). The *lg*1 gene encodes a SQUAMOSA binding protein transactional regulator (Moreno et al., 1997). The *lg1* mutants lack the ligule and auricle at the blade/sheath boundary, which results in more upright leaves (Emerson, 1912; Becraft et al., 1990, Sylvester et al., 1990) and have severely upright leaves.

A metaQTL was identified on chromosome 2 within a ∼1.05 Mb region that was associated with both leaf angle and resistance to maize rough dwarf virus disease (Wang et al., 2016). It was postulated that l*g*1 might be a co-contributor to both the leaf angle (as shown in Tian et al., 2011; Li et al., 2015) and resistance to maize rough dwarf disease (Wang et al., 2016). The expression of *lg1* in leaf tissue (Stelpflug et al., 2016) and the implications of the role of l*g1* in disease resistance found in our study are consistent with a pleiotropic role for this gene. The *Lg1* gene was implicated through GWAS in the nNIL and NAM populations (Poland et al., 2011; Chia et al., 2012), both of which feature B73 as genetic background (approximately 97% and 50%, respectively).

The increases in yield of maize production achieved over the last 50 years has been derived through increased planting population densities, and the indirect selection of corresponding adaptive traits, such as leaf angle (Duvick, 2005) and stress tolerance (Tollenaar and Wu, 1999). In rice, an increase in methyl jasmonate was found to decrease leaf angle by inhibiting brassinosteroid (BR) biosynthesis, the BR signalling pathway, and BR-induced gene expression (Gan et al., 2015). Recently, Tian et al. (2019) dissected a plant architecture QTL that regulates BR and leaf angle, increasing yield in dense population environments, and demonstrated the involvement of LG1 in the regulation of genes controlling these traits. The *sympathy for the ligule* gene, a modifier of the *liguleless narrow1* gene, was also shown to have a pleiotropic role in leaf architecture and disease resistance (Anderson et al., 2019). The observation of both positive and negative correlations between leaf angle and NLB (AUDPC) in the five and four RIL families of the NAM population, respectively, suggests that *lg1* may be multi-allelic, and may have a role in leaves that is distinct from its role in leaf angle and that influences pathogenesis.

The nNIL library characterized in this study is a valuable resource for the genetic characterization and dissection of important traits. The GBS sequencing data provided for this population as a HapMap file allows for highly resolved introgression breakpoints. As with other NIL resources, the population can be utilized to identify and confirm QTL in maize and individual NILs can be used for use in fine-mapping QTL, and for detailed studies on the morphological and physiological mechanisms associated with them. A unique feature of this multi-parental NIL population is its utility for association mapping. Indeed, polymorphisms at *lg1* were identified by association with NLB in this population, confirming the association previously observed in the NAM population by GWAS. The nNIL library can be utilized as a useful resource to further refine traits that have been previously studied in the maize NAM population, or are yet to be characterized.

## Materials and Methods

### NILs

453 backcross five generations, self-pollinated three generations (BC_5_F_3_) NILs derived from the NAM population (Yu et al., 2008) were obtained from the Syngenta Agrochemical Company (Gandhi et al., 2008). The lines were requested with the intention of finding introgressions covering chromosomal segments containing QTL for resistance to NLB, gray leaf spot and aflatoxin accumulation. The lines were chosen by Syngenta based on a proprietary linkage map. The NILs contained introgressions from 18 of donor lines, of which, 13, 3 and 2 donors were derived from the tropical, mixed and NSS sub-populations in maize, respectively (Fig. S2). The lines were selfed in 2010 at the Musgrave Research Farm in Aurora, NY to create BC_5_F_4_ NILs.

### NIL genotyping and analysis

The NIL panel was genotyped by sequencing (Elshire et al., 2011). Four seeds of each line were planted in a 96 cell insert pack and grown under greenhouse conditions. Fresh tissue was harvested from up to 4 seedlings per NIL and DNA was extracted using the Qiagen Plant DNA extraction kit (Qiagen, Germantown, MD, USA). DNA was quantified and checked for quality using the restriction enzyme *EcoR*I. Approximately 30 to 50 ng of DNA was used for 384-plex DNA sequencing at the Institute for Genomic Diversity at Cornell University, Ithaca, NY, USA. SNPs were called using the Tassel5 GBSv1 production pipeline with the ZeaGBSv2.7 TagsOnPhysicalMap (TOPM) file as described in Glaubitz et al. (2014). Imputation was performed with TASSEL-FILLIN (Swarts et al. 2014), using the publicly available haplotype donors file AllZeaGBSv2.7impV5_AnonDonors8k.tar.gz (see https://www.panzea.org/genotypes). GBS data for 955,680 SNPs (B73 AGPv2) were obtained on 412 NILs of the 453 NILs. Of the 41 excluded NILs, 40 had poor or no GBS sequence data and 1 NIL was an exact duplicate of an existing NIL included in the analysis.

### Identification of introgression

Initial graphical genotypes (Young and Tanksley, 1989) were produced for each NIL using an R script that compared GBS-derived DNA sequences and represented each SNP with a vertical line. One set of graphical genotypes compared each NIL with B73. Introgressions were identified by the high density of SNPs relative to B73. In a second analysis, the NIL sequence was compared to its putative introgression donor line. If the donor was correctly identified, the second graphical genotype was the inverse of the first. If the donor was not correctly identified, the introgression would only be seen in the contrast with B73 (Fig. S7).

The introgression parents were further confirmed using the imputed SNP data. For each NIL, all non-B73 SNPs were compared to each of the 18 donor lines. Using Python, this was executed at the whole-genome level, and for each of the putative introgression sites identified (Table S3). If a clear best match was evident for all introgressions and the overall genome, the donor was declared. If several potential donors for a given introgression had similar match scores, the best overall genome match was used to resolve ambiguities (Table S4). The sequence data was also inspected to identify heterozygous regions. The putative donor was also considered in ambiguous cases.

We identified 247 NILs in which the introgressions matched the original putative donor parents based on the graphical genotyping (eg. Fig. S7). Of the remaining lines, 136 and 17 NILs were flagged as putatively incorrect and suspect, respectively. The majority of these cases were readily resolved by comparing SNPs at the specific introgression sites, with one of the donor lines showing a much higher match than the others. In general, the total genome-wide SNP matching was consistent with the interval-specific SNP matching (Table S4). In several cases, however, there was no outstanding donor match; in these cases, inspection of the NIL HapMap file revealed heterozygous introgressions. In these cases, the interval-specific match, genome-wide match and the original putative parent identity were taken into consideration in calling the most likely donor (Table S4). Of the 412 genotyped NILs, 259 (65%) of the NILs matched the original putative parent, and 141 (35%) of the NILs matched another donor parent (Suppl. Data 1). Twelve lines harbored small introgressions that had breakpoints that were determined visually using the HapMap file.

Introgression endpoints were determined using the Python-based program SNPbinner (https://github.com/solgenomics/SNPbinner). SNPbinner was developed for defining breakpoints in GBS data for recombinant inbred lines (Gonda et al., 2018); to our knowledge, this study is the first instance in which this program is being used for NILs. The GBS data was converted to “abh” SNP data in Tassel5 (Bradbury et al., 2007), where ‘a’ was the B73 parent, ‘b’ was the donor parent, and “h” was a heterozygous call. The minimum introgression size was set to 1.5 Mb as a standard for each chromosome. This threshold that was well below 3.25% of the genome size that would be expected for a BC_5_ introgression (ranging from 4.9 to 9.8 Mb per chromosome), but large enough to prevent SNPbinner from breaking apart single introgressions into many small consecutive segments. When necessary, introgressions were visually inspected in Tassel. Nineteen NILs were found to have small introgressions that were not immediately identified via SNP binner and were identified upon visual inspection of the NIL HapMap file in Tassel. The introgressions were visualized using a horizontal line graph in ggplot2 (R Studio) and organized to represent the tiling path across the maize genome.

### Phenotyping NILs and maize diversity panel

The BC_5_F_3_ NILs were grown and self-pollinated at the Cornell University Musgrave Research Farm in Aurora, NY in 2010 to produce BC_5_F_4_ seed. The 453 NILs were planted in randomized complete block designs with two replications per year in 2011, 2012 and 2013 at the same location. The 282 line maize diversity panel (Flint-Garcia et al., 2005) was grown in 2006, 2007, and 2008 in NY and in NC in 2007, as previously reported in Wisser et al. (2011). The diversity panel and the NILs were inoculated with *Setosphaeria turcica* isolate NY001 (race 1) at the 6 to 8 leaf stage in the field as previously described by Chung et al. (2010). Each plant received two types of inoculum: 500 µl of a spore suspension of approximately 4,000 spores/ml and approximately 1.3 ml sorghum kernels that had been colonized by *S. turcica*. The lines were scored three times for DLA after flowering, at intervals of 10 to 14 days. Area under the disease progress curve (Jeger and Viljanen-Rollinson, 2001) per day was determined using the following formula (Das et al., 1992): 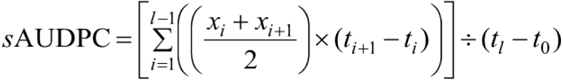, where *x*_*i*_ was the *i*^th^ score and *t*_*i*_ was the *i*^th^ day of measurement, with *i*=1 to *l* measurements.

Diseased leaf area was scored across all replications in each environment for both the NIL and diversity panel trials. Flowering time was scored as DTA for both replications of the NIL trial in 2011 and 2012, and the one replication for the NILs in 2013 as well as in the 282 maize inbred diversity panel. Days to anthesis was determined as the number of days after planting when half of the plants in the row had started pollen shed. The NIL experiment was analyzed as a randomized complete block design in a mixed model including genotype, replication, environment and genotype*environment as random effects to create best linear unbiased prediction (BLUP) estimates for AUDPC and DTA. Variance components were derived to calculate heritability on an entry means basis, where *h*^*2*^ = V_g_/(V_e_/rt+V_gxe_/t+V_g_) (Fehr, 1987).

### Analysis of the NILs for Resistance to NLB

To identify pairwise differences between NILs and the recurrent parent B73, we used a least squares model with AUDPC and DTA as fixed effects to estimate a Dunnett’s pairwise multiple comparison in JMP® Pro Version 13.1.0 (SAS Institute, Inc, Cary, NC, 1989-2019). NILs with AUDPC and DTA values outside the 95% confidence interval of B73 were also identified. The introgressions of the NILs that were significantly more or less resistant to NLB than B73 were compared to prior joint linkage mapping of NLB QTL in the NAM (Poland et al., 2011). These QTL were mapped from germplasm of similar origin (recombinant inbred lines of the NAM), grown at the same location and inoculated with the same NLB isolate. Allelic effects of NLB were estimated for the NILs and compared with the estimates based on the JL QTL in the NAM RIL populations (Suppl Table 2). The QTL designations followed the notation described by Chung et al. (2010), which includes reference to the disease, bin and donor (e.g., qNLB2.01_B73_). If a QTL had multiple founders that contributed resistance, it was designated qNLB2.01_NAM_. A tiling path visualization was created using ggplot2 in R (R Studio), graphically representing the introgressions for each NIL.

### Association mapping in the nNIL library

In order to refine candidate regions and/or genes for resistance to NLB, we endeavored to exploit the sequence differences among the introgression donors, in a uniform B73 genetic background, through association mapping. PROC GLM was utilized in Tassel, using the AUDPC BLUPs for NLB in the nested NIL library. The AGPv2 SNPs were filtered using a minor allele frequency (MAF) of 0.001 and minimum count of 20. Significance thresholds were determined using the Bonferroni correction factor and false discovery rate (FDR). Stepwise regression in Tassel was utilized to identify the SNPs contributing the most towards variation in NLB, using an entry limit of 1 × 10^−6^ and an exit limit of 2 × 10^−6^. Candidate genes and annotated function for the identified SNPs were located using MaizeGDB (www.maizedgb.org; Portland et al., 2018).

### Effect of liguleless1 on NLB

Two *lg1* mutant alleles were tested for resistance to NLB: the *lg1-R* allele (Moreno et al., 1997) in a B73 genetic background, and the *lg1-mum* allele derived from UFMu-1038042 in a W22 genetic background (Settles et al., 2007). The *lg1* mutant lines and corresponding B73 and W22 inbred lines were tested for resistance to NLB in two greenhouse trials, with four replications per trial and 6 to 8 plants per genotype in each replication. Plants were inoculated with a liquid spore suspension in the late afternoon as described above. Overhead sprinklers provided a mist of water for 10 seconds every 10 minutes for approximately 12 to 15 hours. The NLB incubation period was scored as the number of days following inoculation when a necrotic lesion was observed.

### NLB association with leaf angle

Leaf angle measurements for the NAM population were obtained from Panzea (www.panzea.org). Spearman’s correlations were determined using JMP ® Pro Version 13.1.0 between upper leaf angle BLUPs (Tian et al., 2011) and NLB AUDPC BLUPs (Poland et al., 2011) in each of the 26 NAM RIL populations.

### *Genic Association to NLB in* liguleless1

Regional association analysis was conducted using MLM in Tassel with the 282 maize diversity panel based on the NLB AUDPC BLUPs estimated across four environments, with environment, replication nested within environment, and genotype x environment as random effects, and DTA as a fixed effect in BLUP estimation. The HMP3.2.1 LLD SNP dataset that spanned the *lg1* (GRMZM036297) gene was used for this analysis, including 2 kb upstream and downstream of the gene. The MLM analysis used population structure and kinship as previously reported in Samayoa et al. (2015). Bonferonni correction and FDR were used to determine significant SNP associations (Bonferroni et al., 1936; Benjamini and Hochberg, 1995).

## Supporting information

Table S1 nNIL Introgressions

Table S2 nNIL trait data

Table S3 Donor parent matching

Table S4 nNIL donor match compilation

## Acknowledgements

The authors thank various past undergraduate lab members and interns for valuable assistance in the field trials and genotyping the nNIL library, including Alyssa (Cowles) Blachez, Brittany Davenport, Ian Cardle, Nelson Chepkwony, Ariel (Fialko) Chung, Chris Mancuso, Zachary Perry, Theo Pritz, Manuel Romo, and Jacqueline Zee. We thank David Lyon for advice with SNPbinner, and Peter Balint-Kurti, Ed Buckler and Danilo Moreto for helpful comments on the manuscript.

This research was supported by the United States National Science Foundation grant IOS-1127076, USDA-NIFA Hatch under accession# 0227564, USDA-NIFA-AFRI Plant Breeding and Education (Enhancing Education and Research for Plant Disease Resistance (2010-85117-20551)), NIFA Special Projects-Reg Res … Spec Grant: Stories of Crop Evolution, Biodiversity and Domestication, and Methods of Genomic Assisted Crop Improvement for Curricula Development, the McKnight Foundation and Cornell University.

## Supplemental Figures

**Figure S1.**
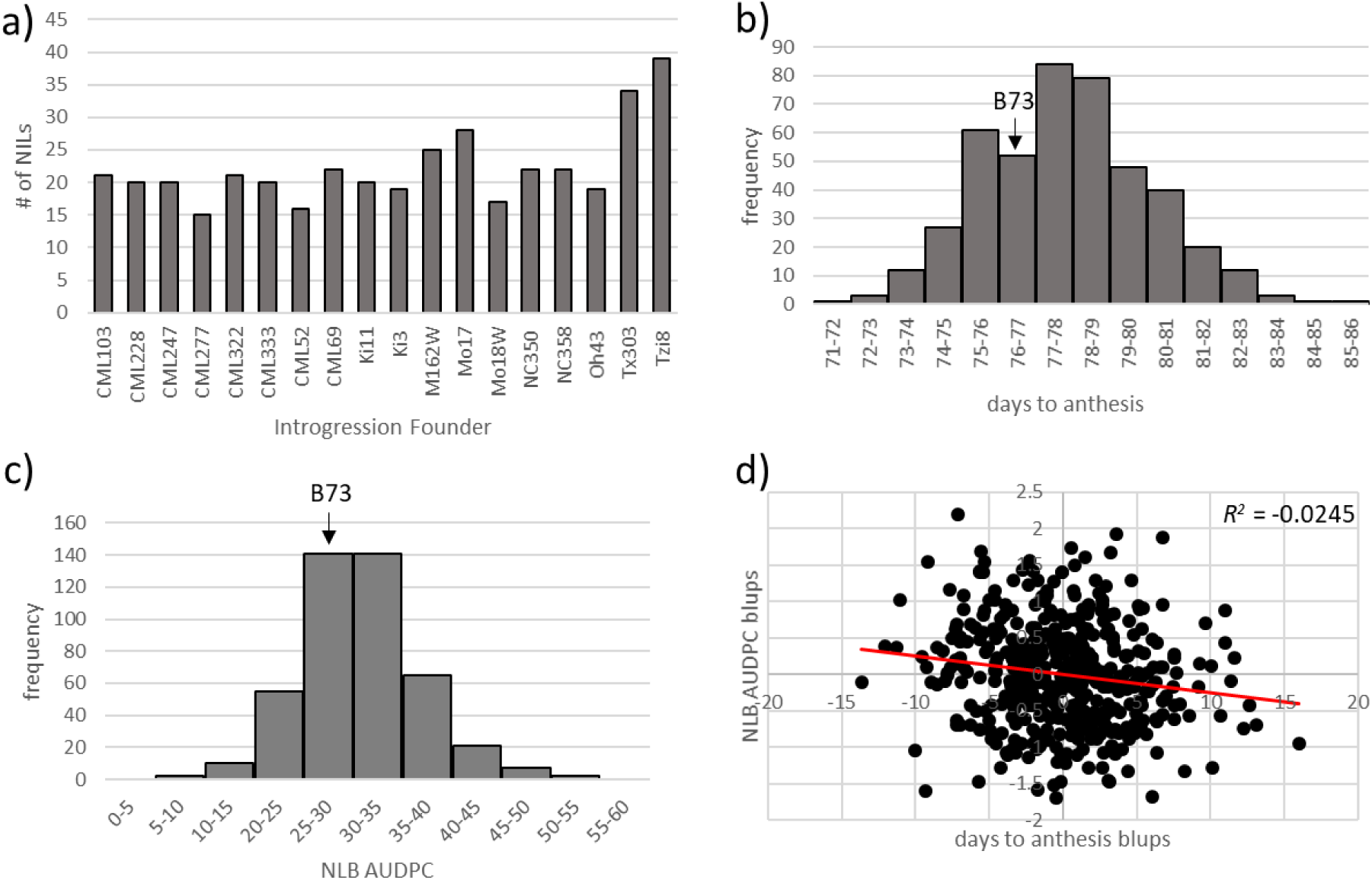
Population metrics in the nNIL library, including a) distribution the number of NILs analyzed in this study based on donor designation from Syngenta, b) frequency distribution of NILs for resistance to NLB, c) frequency distribution of NILs for days to anthesis, and d) genetic correlation of northern leaf blight to days to anthesis. The AUDPC (b) and days to anthesis (c) in the recurrent parent, B73, is shown, indicating that B73 values lie just below the mean of both traits. A small significant genetic correlation (d) was detected between NLB AUDPC and days to anthesis in the nNIL library (c).

**Figure S2.**
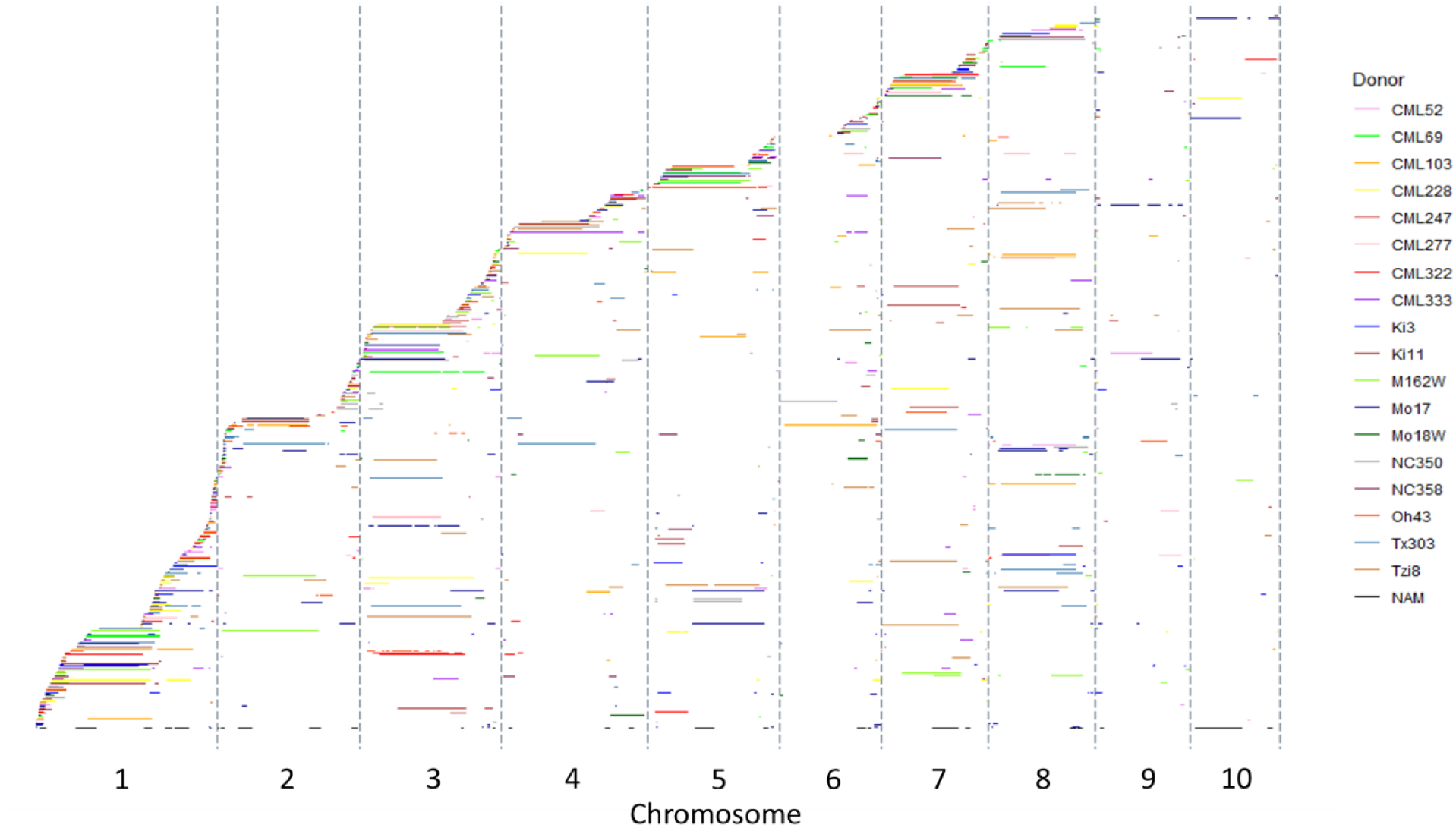
Tiling path of introgressions from nNIL library across the maize genome with introgression donors color-coded to indicate specific donor contributions. NILs were ordered along the y-axis based on the first introgression start site.

**Figure S3.**
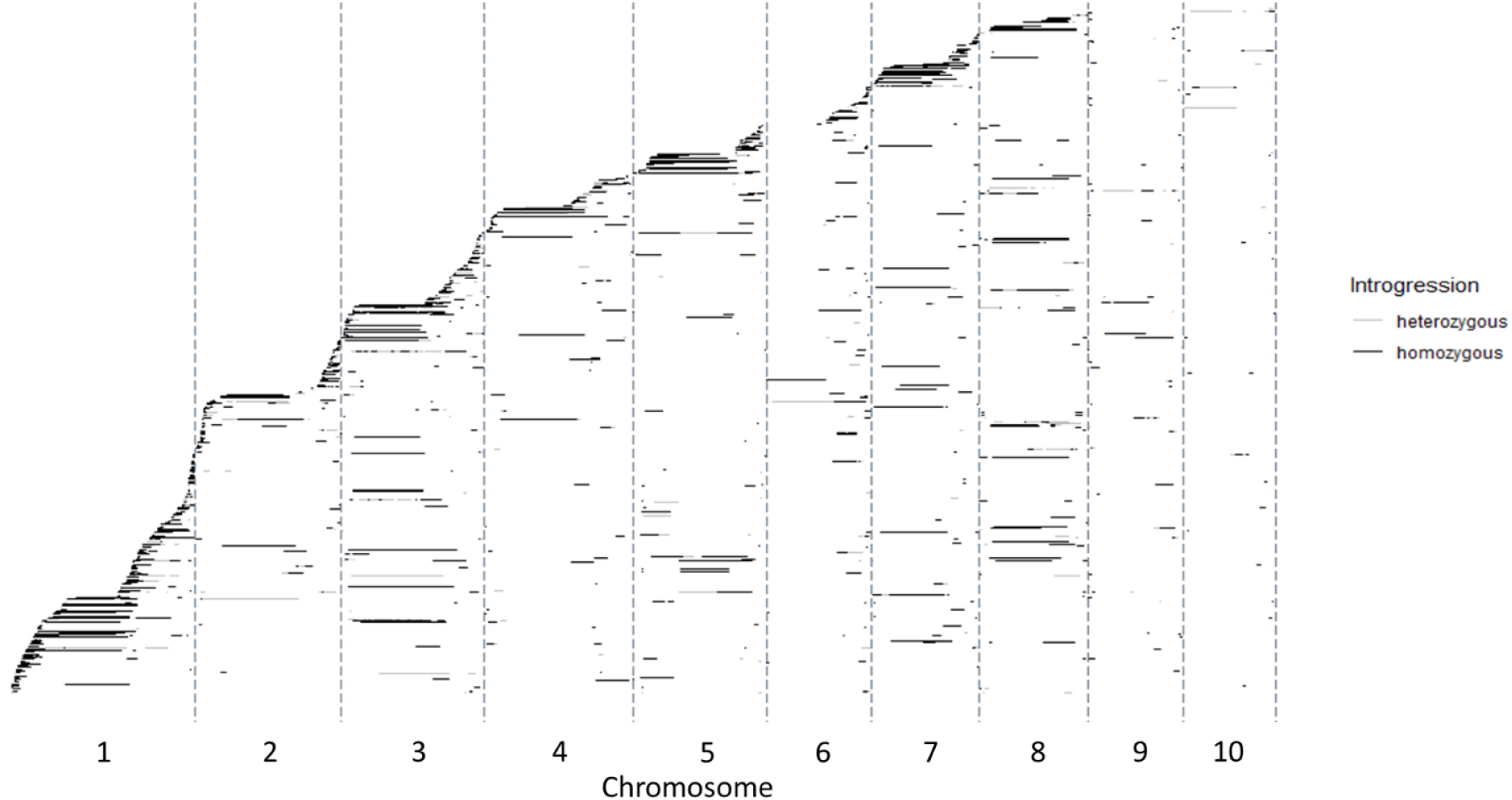
Tiling path of introgressions from nNIL library across the maize genome, with introgressions labelled as either homozygous (black bar) or heterozygous (grey bar). NILs were ordered along the y-axis based on the first introgression start site.

**Figure S4.**
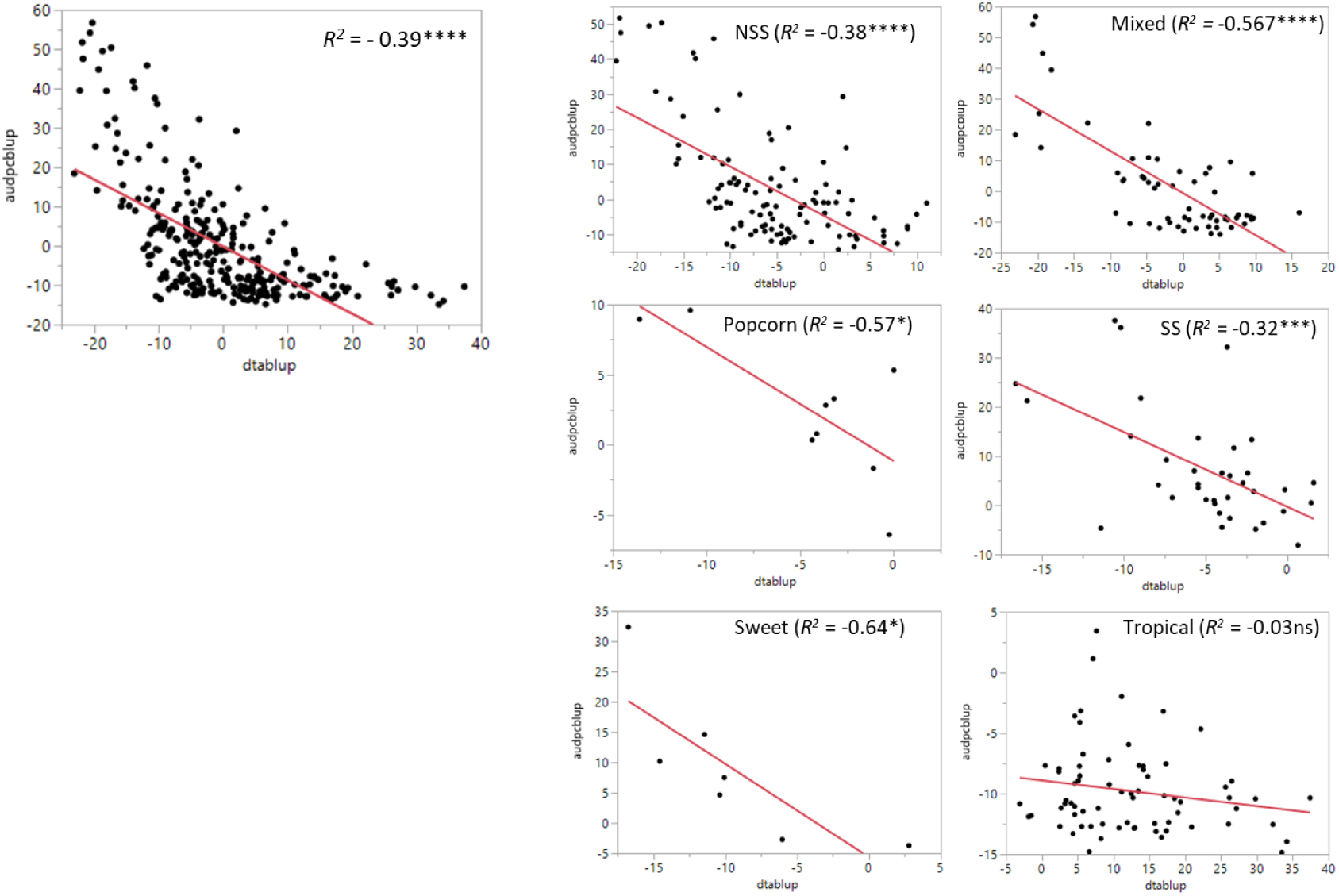
A significant negative correlation was identified between days to anthesis (DTA) and NLB (AUDPC) across the 282 maize diversity panel. The individual subpopulations of the 282 maize diversity population also had significant negative genetic correlations, with the exception of the tropical subpopulation. The tropical subpopulation in general was late flowering with a low level of disease overall.

**Figure S5.**
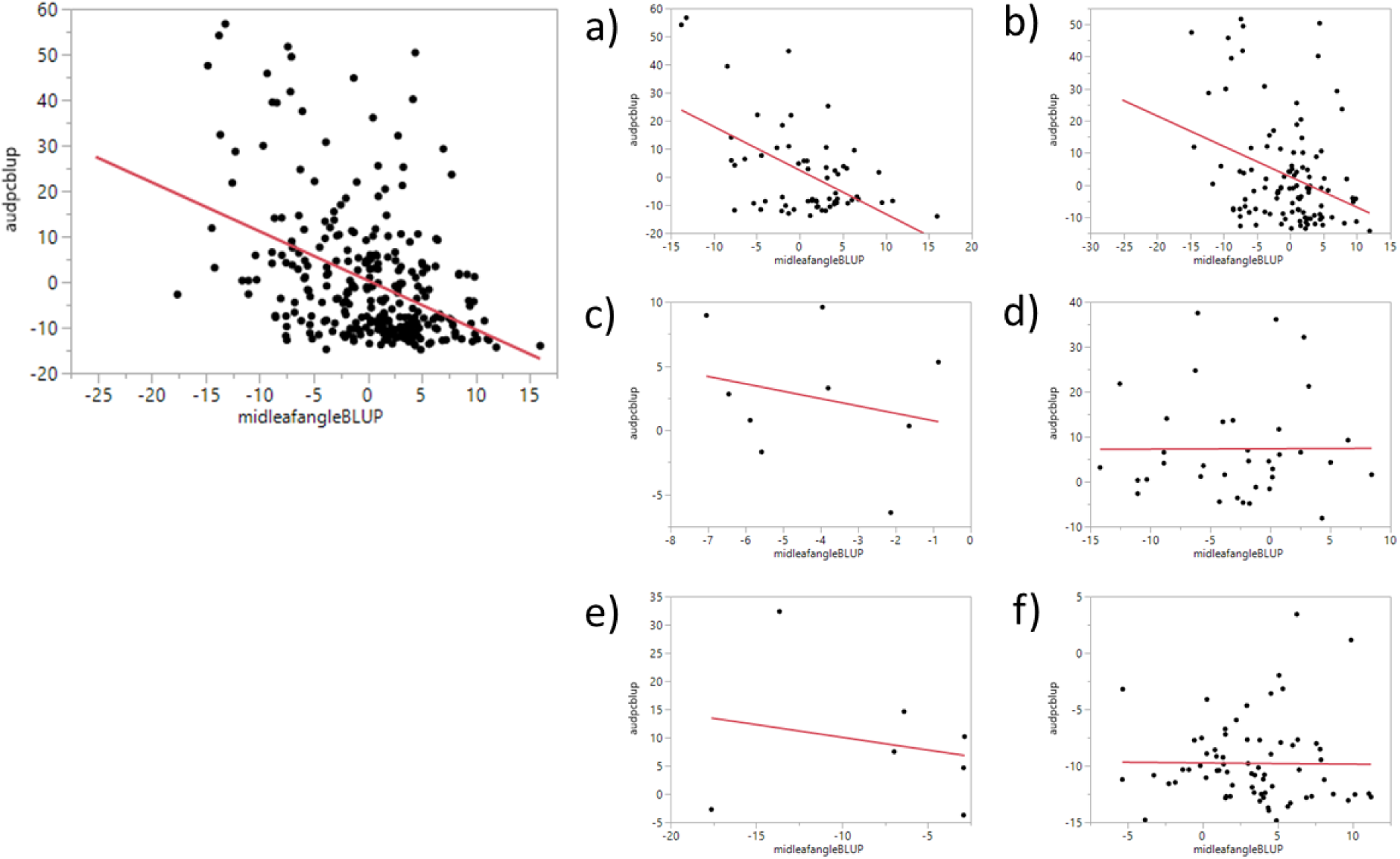
A significant negative correlation was identified between middle leaf angle and NLB (AUDPC) across the 282 maize diversity panel. Significant negative correlations between middle leaf angle and resistance to NLB in the 282 maize inbred diversity panel were identified in the (a) mixed (*p*=0.0001), b) non stiff stalk (*p*=0.006) subpopulations. There was no significant genetic correlation between middle leaf angle and resistance to NLB within the (c) popcorn, (d) stiff stalk, (e) sweet corn, and (f) tropical subpopulations.

**Figure S6.**
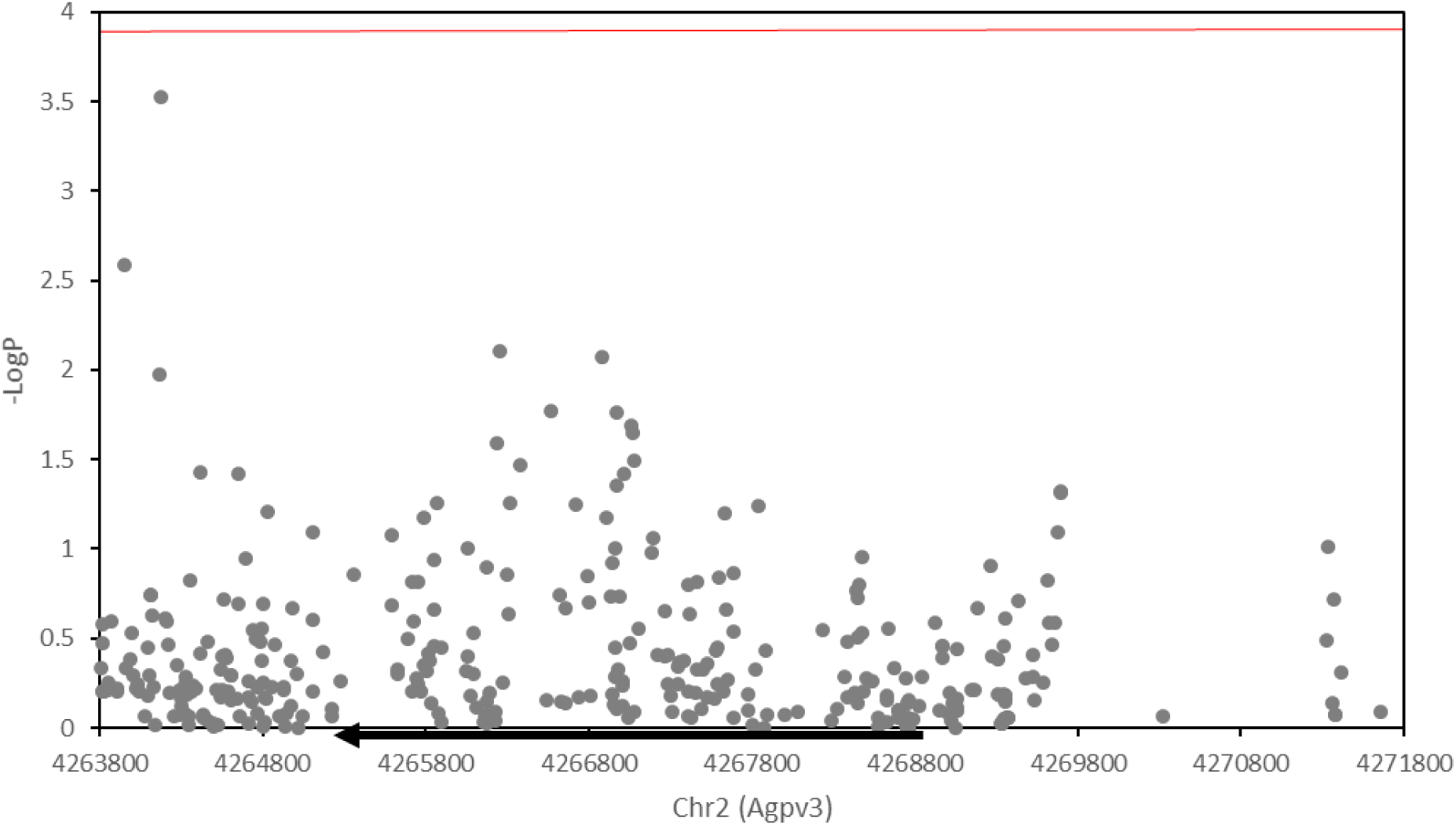
(a) Genic association analysis for resistance to NLB at the *Lg1* genic region using MLM in the 282 maize inbred diversity panel. The black bar along the x-axis represents the location and direction of the *Lg1* gene. No SNPs passed the significant threshold of the Bonferonni correction factor (red line) or the false discovery rate cutoff.

**Figure S7.**
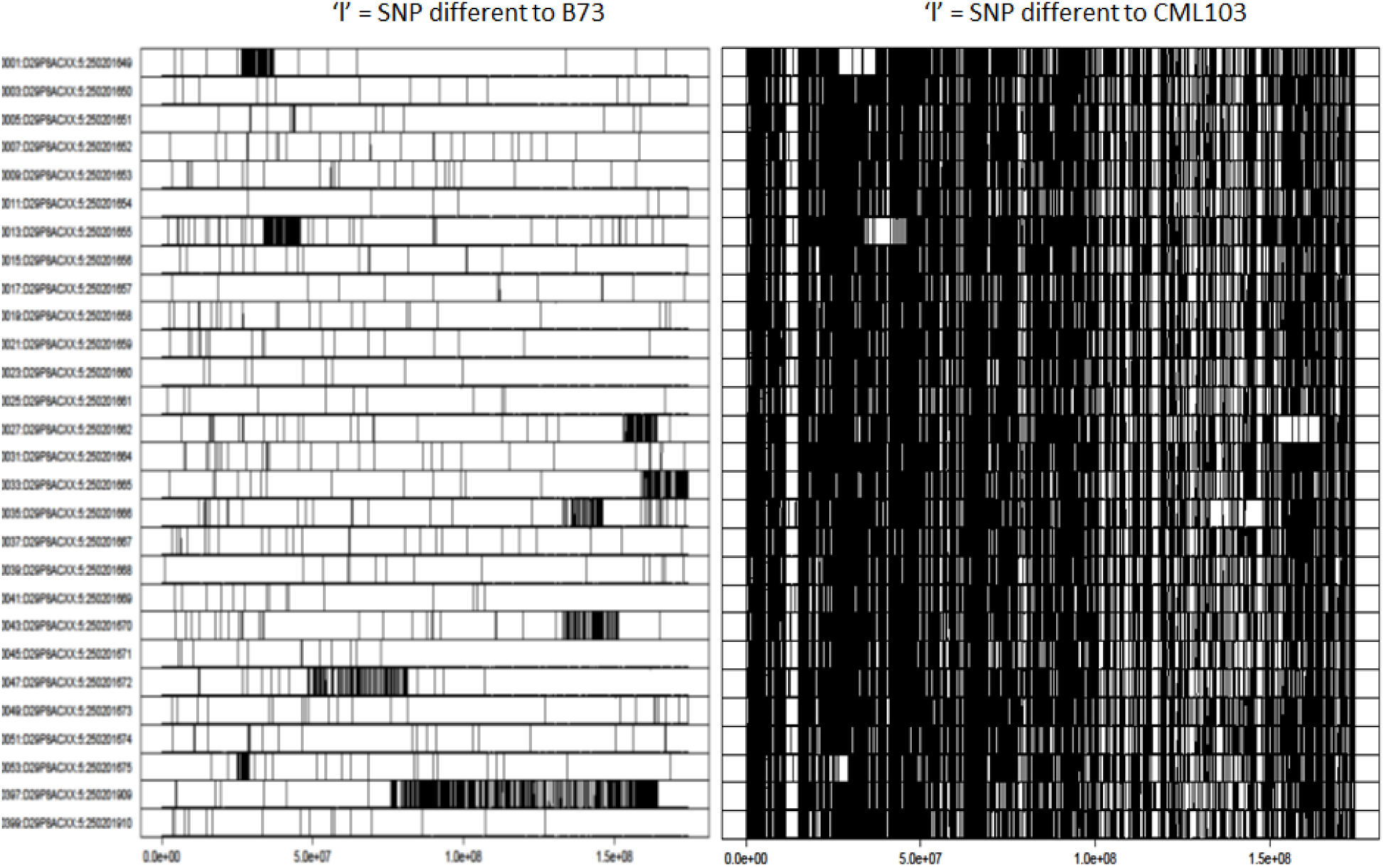
Graphical genotypes of the introgressions along chromosome 6 of the putative CML103-derived NILs. Introgressions are labelled as mismatches, or vertical bars, where the SNP did not match the recurrent parent, B73 (left) and where the SNP did or did not match the putative introgressed parent, CML103 (right). Random bars across the genome implicate GBS sequencing miscalls or mutant SNPs. A concentration of bars indicate a solid introgression region. While several NILs (listed vertically) have corresponding matches between the B73 and CML103 SNP matching, others did not, implicating other introgressed donor(s) for those NILs.

## Supplemental Tables

**Table S1.** NIL data on donor parent designation, introgression start and end points, type of introgression (heterozygous, homozygous or both) and designation with regards to resistance or susceptibility to NLB in comparison to the recurrent parent, B73.

**Table S2.** Least square means for area under the disease progress curve (AUDPC) and days to anthesis (DTA) in the nNIL library. NILs that were significantly different in AUDPC or DTA from B73 in a Dunnett’s test (††), a 95% confidence interval (†), or not significantly different from B73 are noted.

**Table S3.** Parent matching in nNIL library introgressions using SNPs from GBS sequencing to determine the highest SNP match per introgression in comparison to SNPs from donor parent SNPs.

**Table S4.** Verification of the donor parent in a NIL was resolved based on the original parent call, the best introgression SNP match and the best genome SNP match. SNP-matching for donor parents was made only within the 18 donor founder parents used in this study.

## References

Anderson A, et al. (2019) The second site modifier, *Sympathy for the ligule*, encodes a homolog of Arabidopsis ENHANCED DISEASE RESISTANCE4 and rescues the *Liguleless narrow* maize mutant. Plant Cell 31(8):1829–1844.

Balint-Kurti PJ, Yang J, Esbroeck G Van, Jung J, Smith ME (2010) Use of a maize advanced intercross line for mapping of QTL for northern leaf blight resistance and multiple disease resistance. Crop Science 50(2):458–466.

Belcher AR, et al. (2012) Analysis of quantitative disease resistance to southern leaf blight and of multiple disease resistance in maize, using near-isogenic lines. Theor Appl Genet 124(3):433–45.

Benjamini, Y., and Hochberg, Y. (1995) Controlling the false discovery rate: a practical and powerful approach to multiple testing. J. R. Stat. Soc. 57:289–300.

Becraft PW, Bongard-Pierce DK, Sylvester AW, Poethig RS, Freeling M (1990) The *liguleless-1* gene acts tissue specifically in maize leaf development. Dev Biol 141(1):220–232.

Benson JM, Poland JA, Benson BM, Stromberg EL, Nelson RJ (2015) Resistance to gray leaf spot of maize: genetic architecture and mechanisms elucidated through nested association mapping and near-isogenic line analysis. PLoS Genet 11(3):1–23.

Bernacchi D, et al. (1998) Advanced backcross QTL analysis of tomato. II. Evaluation of near-isogenic lines carrying single-donor introgressions for desirable wild QTL-alleles derived from *Lycopersicon hirsutum* and *L. pimpinellifolium*. Theor Appl Genet 97(1–2):170–180.

Bonferroni, C. E., (1936) Teoria statistica delle classi e calcolo delle probabilità, Pubblicazioni del R Istituto Superiore di Scienze Economiche e Commerciali di Firenze.

Bradbury PJ, et al. (2007) TASSEL: Software for association mapping of complex traits in diverse samples. Bioinformatics 23(19):2633–2635.

Brown PJ, et al. (2011) Distinct genetic architectures for male and female inflorescence traits of maize. PLoS Genet 7(11). doi:10.1371/journal.pgen.1002383.

Buckler ES, et al. (2009) The genetic architecture of maize flowering time. Science, 325(5941):714–718.

Chen X, et al. (2010) SQUAMOSA promoter-binding protein-like transcription factors: Star players for plant growth and development. J Integr Plant Biol 52(11):946–951.

Chia J-M, et al. (2012) Maize HapMap2 identifies extant variation from a genome in flux. Nat Genet 44(7):803–7.

Chung CL, Jamann T, Longfellow J, Nelson R (2010) Characterization and fine-mapping of a resistance locus for northern leaf blight in maize bin 8.06. Theor Appl Genet 121(2):205–227.

Chung CL, et al. (2011) Targeted discovery of quantitative trait loci for resistance to northern leaf blight and other diseases of maize. Theor Appl Genet 123(2):307–326.

Cook JP, et al. (2012) Genetic architecture of maize kernel composition in the nested association mapping and inbred association panels. Plant Physiol 158(2):824–834.

Das MK, Rajaram S, Mundt CC, Kronstad WE (1992) Inheritance of slow-rusting resistance to leaf rust in wheat. Crop Sci 32(6):1452.

Ding X, et al. (2017) Both major and minor QTL associated with plant height can be identified using near-isogenic lines in maize. Euphytica 213(1). doi:10.1007/s10681-016-1825-9.

Ducrocq S, et al. (2008) Key impact of *Vgt1* on flowering time adaptation in maize: Evidence from association mapping and ecogeographical information. Genetics 178(4):2433–2437.

Duvick DN (2005) The Contribution of Breeding to Yield Advances in maize (Zea mays L.). Adv Agron 86:83–145.

Dzievit MJ, Li X, Yu J (2019) Dissection of leaf angle variation in maize through genetic mapping and meta-analysis. Plant Genome 12(1):1–12.

Elshire RJ, et al. (2011) A robust, simple genotyping-by-sequencing (GBS) approach for high diversity species. PLoS One 6(5):1–10.

Emerson, RA (1912). The inheritance of the ligule and auricles of corn leaves. Nebraska Agric. Expt. Stn. Annu. Rep. 25:81–88.

Eshed Y, Zamir D (1995) An introgression line population of *Lycopersicon pennellii* in the cultivated tomato enables the identification and fine mapping of yield-associated QTL. Genetics 141(3):1147–1162.

Fehr WR (1987) Principles of cultivar development: theory and technique, Vol 1. Macmillan, New York

Flint-Garcia SA, et al. (2005) Maize association population: A high-resolution platform for quantitative trait locus dissection. Plant J 44(6):1054–1064.

Galiano-Carneiro AL, Miedaner T (2017) Genetics of resistance and pathogenicity in the maize/*Setosphaeria turcica* pathosystem and implications for breeding. Front Plant Sci 8(August):1–13.

Gan L, et al. (2015) Methyl jasmonate inhibits lamina joint inclination by repressing brassinosteroid biosynthesis and signaling in rice. Plant Sci 241:238–245.

Gandhi, S., et al. (2008). High throughput allele discovery and incorporation in elite maize germplasm. Proc for the 5th International Crop Science Congress.

Glaubitz JC, et al. (2014) TASSEL-GBS: A high capacity genotyping by sequencing analysis pipeline. PLoS One 9(2). doi:10.1371/journal.pone.0090346.

Gonda I, et al. (2019) Sequencing-based bin map construction of a tomato mapping population, facilitating high-resolution quantitative trait loci detection. Plant Genome 12(1):1–14.

Hung H-Y, et al. (2012) ZmCCT and the genetic basis of day-length adaptation underlying the postdomestication spread of maize. Proc Natl Acad Sci 109(28):E1913–E1921.

Hurni S, et al. (2015) The maize disease resistance gene *Htn1* against northern corn leaf blight encodes a wall-associated receptor-like kinase. Proc Natl Acad Sci 112(28):8780–8785.

Jamann TM, et al. (2016) A remorin gene is implicated in quantitative disease resistance in maize. Theor Appl Genet 129(3):591–602.

Jamann TM, Poland JA, Kolkman JM, Smith LG, Nelson RJ (2014) Unraveling genomic complexity at a quantitative Disease resistance locus in maize. Genetics 198(1):333–344.

Jeger MJ, Viljanen-Rollinson SLH (2001) The use of the area under the disease-progress curve (AUDPC) to assess quantitative disease resistance in crop cultivars. Theor Appl Genet 102(1):32–40.

Jeuken MJW, Lindhout P (2004) The development of lettuce backcross inbred lines (BILs) for exploitation of the *Lactuca saligna* (wild lettuce) germplasm. Theor Appl Genet 109(2):394–401.

Johnston R, et al. (2014) Transcriptomic analyses indicate that maize ligule development recapitulates gene expression patterns that occur during lateral organ initiation. Plant Cell Online 26(12):4718–4732.

Keurentjes JJB, et al. (2007) Development of a near-isogenic line population of *Arabidopsis thaliana* and comparison of mapping power with a recombinant inbred line population. Genetics 175(2):891–905.

Kump KL, et al. (2011) Genome-wide association study of quantitative resistance to southern leaf blight in the maize nested association mapping population. Nat Genet 43(2):163–168.

Liang Z, Schnable JC (2016) RNA-seq based analysis of population structure within the maize inbred B73. PLoS One 11(6):1–15.

Li C, et al. (2015) Genetic control of the leaf angle and leaf orientation value as revealed by ultra-high density maps in three connected maize populations. PLoS One 10(3):1–13.

Li Y, et al. (2018) Increased experimental conditions and marker densities identified more genetic loci associated with southern and northern leaf blight resistance in maize. (April):1–12.

Lopez-Zuniga LO, et al. (2019) Using maize chromosome segment substitution line populations for the identification of loci associated with multiple disease resistance. G3 Genes, Genomes, Genet 9(1):189–201.

Mackill DJ, Bonman JM (1992) Inheritance of blast resistance in near-isogenic lines of rice. Phytopathology 82 (7): 746–749.

Marchadier E, et al. (2019) The complex genetic architecture of shoot growth natural variation in Arabidopsis thaliana. PLoS Genet 15(4):e1007954.

McMullen MD, et al. (2009) Genetic properties of the maize nested association mapping population. Science, 325(5941), 737–740.

Monforte AJ, Tanksley SD (2000) Development of a set of near isogenic and backcross recombinant inbred lines containing most of the *Lycopersicon hirsutum* genome in a *L. esculentum* genetic background: A tool for gene mapping and gene discovery. Genome 43(5):803–813.

Moreno MA, Harper LC, Krueger RW, Dellaporta SL, Freeling M (1997) *Liguleless1* Encodes a Nuclear-Localized Protein Required for Induction of Ligules and Auricles During Maize Leaf Organogenesis. Genes Dev 11(5):616–628.

Olukolu BA, et al. (2014) A genome-wide association study of the maize hypersensitive defense response identifies genes that cluster in related pathways. PLoS Genet 10(8):e1004562.

Peiffer JA, et al. (2013) The Genetic Architecture of Maize Stalk Strength. PLoS One 8(6). doi:10.1371/journal.pone.0067066.

Peiffer JA, et al. (2014) The genetic architecture of maize height. Genetics 196(4):1337–1356.

Poland JA, Bradbury PJ, Buckler ES, Nelson RJ (2011) Genome-wide nested association mapping of quantitative resistance to northern leaf blight in maize. Proc Natl Acad Sci U S A 108(17):6893–8.

Portwood JL, et al. (2019) MaizeGDB 2018: The maize multi-genome genetics and genomics database. Nucleic Acids Res 47(D1):D1146–D1154. doi: 10.1093/nar/gky1046

Rodgers-Melnick E, et al. (2015) Recombination in diverse maize is stable, predictable, and associated with genetic load. Proc Natl Acad Sci 112(12):201413864.

Salvi S, et al. (2007) Conserved noncoding genomic sequences associated with a flowering-time quantitative trait locus in maize. Proc Natl Acad Sci U S A 104(27):11376–11381.

Samayoa L, Malvar R, Olukolu BA, Holland JB, Butrón A (2015) Genome-wide association study reveals a set of genes associated with resistance to the Mediterranean corn borer (*Sesamia nonagrioides* L.) in a maize diversity panel. BMC Plant Biol 15(1):35.

Settles AM, et al. (2007) Sequence-indexed mutations in maize using the UniformMu transposon-tagging population. BMC Genomics 8:1–12.

Springer NM, et al. (2009) Maize inbreds exhibit high levels of copy number variation (CNV) and presence/absence variation (PAV) in genome content. PLoS Genet 5(11). doi:10.1371/journal.pgen.1000734.

Stelpflug SC, et al. (2016) An expanded maize gene expression atlas based on RNA Sequencing and its use to explore root development. Plant Genome 9(1):0.

Swarts K, et al. (2014) Novel methods to optimize genotypic imputation for low-coverage, next-generation sequence data in crop plants. Plant Genome 7(3):0.

Sylvester a W, Cande WZ, Freeling M (1990) Division and differentiation during normal and *liguleless-1* maize leaf development. Development 110(3):985–1000.

Tian F, et al. (2011) Genome-wide association study of leaf architecture in the maize nested association mapping population. Nat Genet 43(2):159–162.

Tian J, et al. (2019) Teosinte ligule allele narrows plant architecture and enhances high-density maize yields. Science (80-) 365(6454):658–664.

Tollenaar M, Wu J (1999) Yield improvement in temperate maize is attributable to greater stress tolerance. Crop Sci 39(6):1597–1604.

Tuinstra MR, Ejeta G, Goldsbrough PB (1997) Heterogeneous inbred family (HIF) analysis?: a method for developing near-isogenic lines that differ at quantitative trait loci. Theor Appl Genet 95(5-6):1005–1011.

Wallace JG, et al. (2014) Association mapping across numerous traits reveals patterns of functional variation in maize. PLoS Genet 10(12). doi:10.1371/journal.pgen.1004845.

Wang Y, et al. (2016) A comprehensive meta-analysis of plant morphology, yield, stay-green, and virus disease resistance QTL in maize (Zea mays L.). Planta 243(2):459–471.

Welz HG, Geiger HH (2000) Genes for resistance to northern corn leaf blight in diverse maize populations. Plant Breed 119(1):1–14.

Wisser RJ, Balint-Kurti PJ, Nelson RJ (2006) The genetic architecture of disease resistance in maize: a synthesis of published studies. Phytopathology 96(2):120–129.

Wisser RJ, et al. (2011) Multivariate analysis of maize disease resistances suggests a pleiotropic genetic basis and implicates a GST gene. Proc Natl Acad Sci U S A 108(18):7339–7344.

Yang Q, et al. (2017) A gene encoding maize caffeoyl-CoA O-methyltransferase confers quantitative resistance to multiple pathogens. Nat Genet 49:1364.

Young ND, Zamir D, Ganal MW, Tanksley SD (1988) Use of isogenic lines and simultaneous probing to identify DNA markers tightly linked to the *Tm-2a* gene in tomato. Genetics 120(2):579–586.

Young ND, Tanksley SD (1989) Restriction fragment length polymorphism maps and the concept of graphical genotypes. Theor Appl Genet 77(1):95–101.

Yu J, Holland JB, McMullen MD, Buckler ES (2008) Genetic design and statistical power of nested association mapping in maize. Genetics 178(1):539–551.

Zhang X, Yang Q, Rucker E, Thomason W, Balint-Kurti P (2017) Fine mapping of a quantitative resistance gene for gray leaf spot of maize (Zea mays L.) derived from teosinte (*Z. mays* ssp. *parviglumis*). Theor Appl Genet 130(6):1285–1295.

